# BirT: a novel primer pair for avian environmental DNA metabarcoding

**DOI:** 10.1101/2023.08.08.552521

**Authors:** B. Thalinger, R Empey, M. Cowperthwaite, K Coveny, D. Steinke

## Abstract

Environmental DNA metabarcoding has become a widely used technique to detect animals from environmental samples and is on the brink of being implemented into routine species monitoring. Surprisingly, birds are among the taxonomic groups which until recently, received comparably little attention in eDNA research, a fact that is changing rapidly, with the growing number of air eDNA analyses. Since birds are hardly ever the most abundant species in aquatic or terrestrial habitats, a high specificity of the employed metabarcoding primers is key to limit non-target amplifications and enable reliable detection from environmental samples.

Here, we present a novel primer pair (BirT) for metabarcoding of avian environmental DNA. We optimized specificity and fragment length regarding taxonomic resolution and available sequencing technology. Additionally, we evaluated the availability of 12S reference sequences for birds and filled database gaps by generating novel 12S barcodes. Finally, we tested the applicability of this approach using field-collected eDNA samples obtained with three different filter types. These results were compared to visual observations uploaded to eBird (www.eBird.org) during the sampling period.

Our results confirm the suitability of the BirT primer pair for bird eDNA metabarcoding with optimized fragment length, no amplification of key non-target groups, and taxonomic resolution provided by the amplified fragment. Albeit there are still substantial gaps in the 12S reference sequence database, the analysis of bird eDNA from water samples resulted in species-level taxonomic resolution for 92% of the detected taxa. All tested filter/filtration combinations delivered similar results for total read numbers per sample (mean: 613,972 ± 340,088 SD) and species detected per sample (mean: 5.5 ± 2.3 SD). Ninety-five percent of the bird detections were highly plausible and 58% confirmed by visual observations. The majority of the detected bird species was closely associated with aquatic habitats confirming the suitability of water samples for the detection of waterfowl and species inhabiting similar ecological niches via eDNA.

## Introduction

Environmental DNA metabarcoding has become a widely used technique to detect animals from environmental samples and is on the brink of being implemented into routine species monitoring (Bruce et al., 2021; Yates, Furlan, Thalinger, Yamanaka, & Bernatchez, 2023). This approach allows distinguishing between species based on differences between DNA barcodes generated via high throughput sequencing which are subsequently mapped to reference sequence databases for taxonomic identification (Hebert, Cywinska, Ball, & DeWaard, 2003; Yu et al., 2012). The requirement of metabarcoding primers to be specific to one or several target taxa and the conditions under which mismatches at the priming site can be tolerated have received much attention in the scientific discourse. On one hand, primer pairs specific to a certain taxon (e.g. Aves), which at the same time exhibit mismatches at the priming sites for other taxa, will deliver sequencing results primarily associated with the target taxon, albeit at low quality in case only little or degraded target DNA is present in the sample (Elbrecht & Leese, 2017b; Josep Piñol, Senar, & Symondson, 2019). On the other hand, primer pairs which are more “universal” and capable of amplifying eDNA from a group of taxa, (e.g. all vertebrates (Riaz et al., 2011)) likely detect a higher number of locally occurring species without additional sequencing efforts (Mariani et al., 2021) at the cost of losing sequencing depth to taxa not originally under investigation (Collins et al., 2019; Klure, Greenhalgh, & Dearing, 2022; Levesque-Beaudin, Steinke, Böcker, & Thalinger, 2023; Yang, Zhang, Mu, & Zhang, 2023). Additionally, universal primers are more prone to lower amplification efficiency due to mismatches at the priming site (Elbrecht & Leese, 2015; J. Piñol, Mir, Gomez-Polo, & Agustí, 2015; Bista et al., 2018) which can only be partly compensated using degenerate bases in primers (Elbrecht et al., 2019; Tournayre et al., 2020). To date, universal primers are mostly employed for metabarcoding especially if not much is known about the species occurring in an environment or if the detection of a broad range of taxa with minimum laboratory and sequencing efforts is required (Holman et al., 2021; Ficetola & Taberlet, 2023). However, a growing number of optimized eDNA metabarcoding primers have been designed for more targeted research questions, existing primer pairs were modified (e.g. Valsecchi et al., 2020; Min, Barber, & Gold, 2021; Newton et al., 2023), and several primer pairs used in combination to expand taxonomic coverage (Stat et al., 2017; Li, Qin, Wang, Zhang, & Yang, 2023).

Until recently, Aves were one of the taxonomic groups receiving little attention in eDNA research (Thalinger, Deiner, et al., 2021; Takahashi et al., 2023). The reasons for this are twofold. Visual observation is still the most frequent used method of detection despite some shortcomings (Kamp, Oppel, Heldbjerg, Nyegaard, & Donald, 2016) and the recent advent of acoustic sensing methods supported by AI caught a lot of attention (Aide et al., 2013). In some studies, bird species appeared as additional detections in fish eDNA metabarcoding datasets (Mariani et al., 2021; Jensen et al., 2023) or were part of eDNA vertebrate community surveys (Johnson, Barnes, Garrett, & Clare, 2023; Zhang, Zhao, & Yao, 2023). More recently, Ushio et al. (2018) and Taberlet et al. (2018) developed primer pairs specifically for bird eDNA metabarcoding. However, technological advances in combination with the boom of air environmental DNA analysis (e.g. Littlefair et al., 2023; Lynggaard, Frøslev, Johnson, Olsen, & Bohmann, 2023; Bohmann & Lynggaard, 2023) warrant a re-evaluation of the available primer sets for bird eDNA metabarcoding and the conditions under which this approach could be applicable in future large-scale field studies.

A high quality approach for eDNA metabarcoding requires a) a fragment length suitable for the detection of environmental DNA (Jo, Takao, & Minamoto, 2022) and for most high throughput sequencing platforms with bi-directional approaches (Deiner et al., 2017), b) a target fragment encompassing areas of high interspecific sequence variability but limited intraspecific variability (Jackman et al., 2021), c) the availability of a reference sequence database enabling taxonomic identification (Weigand et al., 2019; Jackman et al., 2021), and d) a target gene which fulfills all previously mentioned conditions and occurs in high copy numbers (for this reason mitochondrial genes are predominantly used for the detection of animal eDNA) (Jo et al., 2022).

A fragment length of 80 to 400 bp was originally deemed appropriate for the detection of trace amounts of degraded DNA (King, Read, Traugott, & Symondson, 2008). Although there are many case studies indicating a higher abundance of short DNA fragments in the environment, others showed the prevalence of high-quality, long fragments of eDNA (Thalinger, Deiner, et al., 2021; Jo et al., 2022) rendering short target fragments not the highest priority for successful detection of target species. Additionally, sequencing technologies are constantly advancing with the increased availability of long-range sequencing (Ritter et al., 2020) and the availability of high throughput sequencing kits running for 500 or 600 cycles. The latter allow the amplification of target fragments (incl. all primers, adaptors, tags,…) slightly below 500 and 600 bp, respectively, and are available for instruments producing more than 500mio paired end reads per run (Deiner et al., 2017; McClenaghan et al., 2020).

A successful species-level match of sequence reads to reference barcodes requires closely related, co-occurring species to exhibit different DNA barcodes (Deiner et al., 2017; Jackman et al., 2021). Short fragments below 100 bp and including only a single area of high sequence variability have been deemed suitable for such taxonomic assignments in environments that contain only a limited number of target taxa (e.g. Taberlet et al., 2018). However, usually longer fragments between 180 and 450 bp are used for eDNA metabarcoding of more complex communities because the number of highly variable sites and the chances of finding distinct reference barcodes for each species increase with fragment length (Zhang, Zhao, & Yao, 2020).

The mitochondrial 12S rRNA (12S) gene is frequently used for the detection of vertebrates in environmental samples (Sales et al., 2020; Zhang et al., 2020). Most available primer pairs are less prone to co-amplify microorganisms, fungi, arthropods, and other non-vertebrate groups (Deagle, Jarman, Coissac, Pompanon, & Taberlet, 2014; Collins et al., 2019). However, previous vertebrate barcoding efforts have focused on the Cytochrome B and Cytochrome Oxidase I genes and thus no large-scale, international efforts were ever made to obtain 12S reference sequences for all vertebrates (Jackman et al., 2021). For birds, previous attempts at primer design for metabarcoding used a limited amount of publicly available sequences or other target genes (Ushio et al., 2018; Newton et al., 2023).

Here we present a novel primer pair for metabarcoding avian environmental DNA. We optimized the specificity of the primers to bird species and the fragment length with to desired taxonomic resolution and available sequencing technology. Additionally, we evaluated the availability of 12S reference sequences for birds and tried to close some of the database gaps by generating novel 12S barcodes. Finally, we tested the applicability of this approach with field-collected eDNA samples using three different filter types and by comparing the obtained results to visual observations uploaded to eBird (www.eBird.org) during the sampling period.

## Materials and Methods

### Available reference sequences and primer design

To assess the taxonomic coverage of available 12S references sequences for birds, Genbank was queried (originally on 1^st^ Oct 2019; a final time on 16^th^ Jan 2023) for all available sequences of the mitochondrial 12S rRNA (12S) gene associated with “Aves”. Out of the 9,821 downloaded sequences, 2,828 stemming from population genomics work were removed (SI 1), except for a few sequences per species. Only sequences actually containing the 12S region were extracted leading to the removal of another 101 sequences (SI 1). The remaining 6,889 sequences were aligned using the MAFFT FFT-NS-1 algorithm (MAFFT v7.388) in Geneious Prime® (version 2022.0.1). Several primer pairs previously developed for 12S bird DNA metabarcoding were added to the alignment (Fig. 1) and the 2,621 sequences covering the entire length amplified by these primers as well as sequences of selected non-target taxa (SI 1 and Fig. 2) were used to manually evaluate the potential of individual priming sites and primer combinations. Our aim was to design primers with similar melting temperatures of approx. 57°C. Additionally, we aimed for minimal effects of secondary structures. Specifics of the potential primers were checked with the OligoAnalyzer^TM^ Tool (Integrated DNA Technologies). The optimal primers should have sufficient mismatches with non-target taxa such as fish, amphibians, and humans to at least restrict their amplification. At the same time, we tried to choose priming sites for which the base-code is as similar as possible for all bird species, to avoid the use of degenerate bases as much as possible (Miya et al., 2015; Elbrecht et al., 2019). Additionally, the selected fragment should deliver a high taxonomic resolution and allow to distinguish between individual bird species, while at the same time being short enough to amplify degraded DNA and have sufficient overlap between forward and reverse reads on Illumina sequencing platforms operated with 500 or 600 cycle chemistry. All primer design work was carried out by researchers with more than a decade of experience, hence, forgoing automated solutions for primer design such as PrimerMiner (Elbrecht & Leese, 2017a), PrimerQuest^TM^ (Integrated DNA Technologies), or Primer 3 (Untergasser et al., 2012).

**Figure 1:**
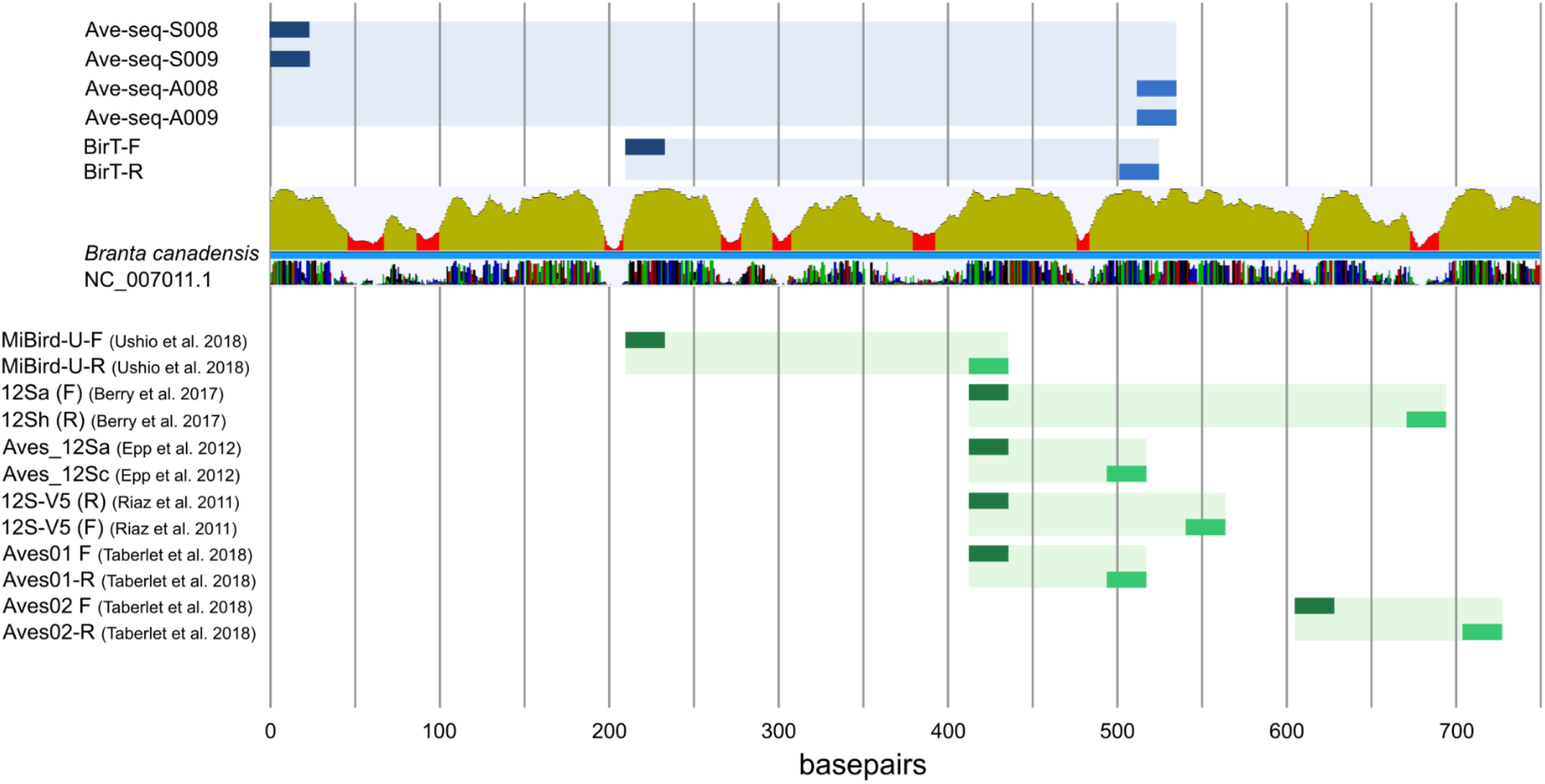
The top half of the figure displays the position of the developed Sanger sequencing primers (Ave-seq) and the BirT primer pair developed for bird eDNA metabarcoding. Dark blue boxes show the position of forward primers, bright blue boxes the position of reverse primers, and the light blue boxes connecting them depict the amplified fragment in relation to the 12S gene of *Branta canadensis* (accession number NC_007011.1). Based on the analysis of all available 12S bird sequences, red areas indicate high sequence variability, while peaks indicate areas of low sequence variability, which are suitable as priming sites for general bird primers. The bottom half of the figure displays the position and amplified fragment of previously published 12S primers used for the analysis of eDNA. Forward primers shown in dark green, reverse primers in bright green. The connecting light green represents the amplified fragment.

**Figure 2:**
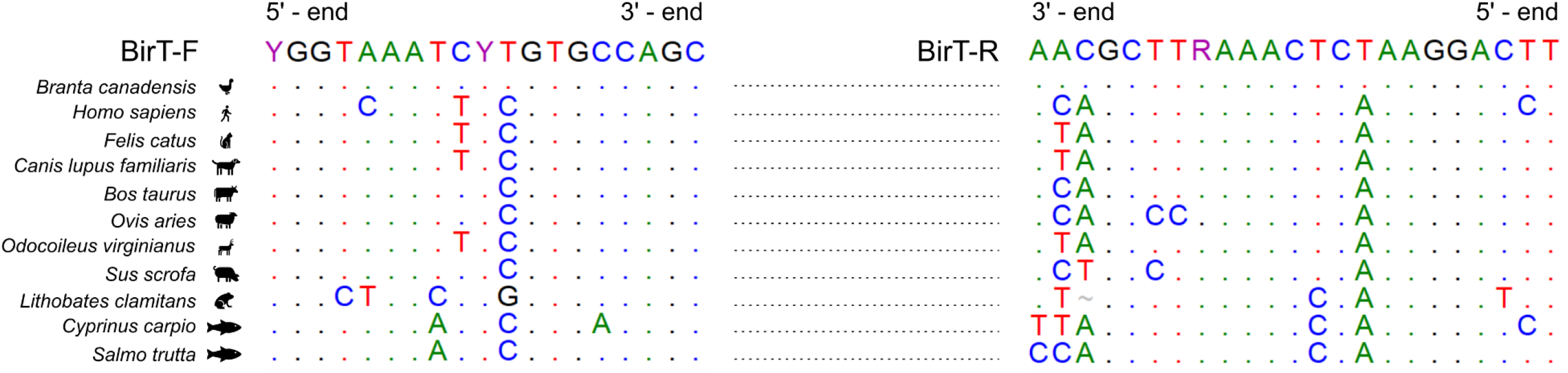
Alignment of BirT primers with important non-target organisms at both forward and reverse priming sites. Identical bases in relation to the primers are displayed as dots, while the basecode is provided for all mismatches. Please note that the primers are displayed to reflect their position and direction in an alignment (i.e. the BirT-R primer is displayed as reverse complement) and to facilitate the display of the mismatches and their location with respect to the 3’ and 5’ ends.

After the selection of a primer-pair, available reference sequences were re-evaluated for the fragment in question. Of the previously downloaded reference sequences, 5472 spanned the entire selected fragment. The taxonomy of these sequences was compared to the IOC World Bird List (v 12.2) (Donsker & Rasmussen, 2022) which contains 9,862 entries of non-extinct species. Using Avibase (https://avibase.bsc-eoc.org/), the species without a match were manually checked for changes in nomenclature, being birds, or being extinct. Affected sequences were renamed or removed from the database. Finally, the sequences were aligned to a *Branta canadensis* reference sequence (GenBank Accession number NC_007011.1) and all sequences which could not be aligned were discarded resulting in 4,544 remaining sequences. Their taxonomic distribution with respect to the entire bird taxonomy was plotted to reveal gaps in the reference database.

### Reduction of reference sequence library gaps

To reduce the substantial gaps in the reference sequence database starting with “locally occurring” species, a list of Canadian bird species was compiled from Avibase and compared to the already available 12S sequences. They were sorted based on their association with water by an expert from Bird Studies Canada (D. Tozer). Thereafter, bird species, which closely associate with aquatic habitats and without previously available 12S sequences were given priority for the generation of new reference sequences from tissue samples.

First, the DNA extracts and tissue samples available at the Centre for Biodiversity Genomics (University of Guelph, Canada) were compiled. Next, tissue samples were obtained from the reference collection at the Royal Ontario Museum (Toronto, Canada). We aimed at sub-sampling three samples from separate individuals for each bird species and if possible, incorporated samples from different subspecies. Whenever feasible, these efforts also encompassed species with distribution ranges outside of Canada. Tissues were subsampled using forceps and scalpels either dipped in 96% ethanol and singed three times or cleaned with Eliminase^®^(VWR, Canada) and nuclease-free water, subsequently. The tissue samples were transported to the University of Guelph in a cooling box and stored at −20°C until further processing. All subsequent laboratory work was carried out in designated rooms and included the strict separation of pre-PCR and post-PCR workflows, the use of UV-ed, hooded workspaces, bleach-cleaned laboratory surfaces and the wearing of protective clothing and DNA-free gloves at all times. Tissues containing ethanol as a means of preservation were dried in a fume hood. The DNeasy Blood and Tissue Kit (Qiagen) was used for lysis and extraction: 200 µl lysis buffer consisting of 180 µl ATL buffer and 20 µl Proteinase K (20 mg/ml) were added to each tissue followed by incubation over night at 56°C on a rocking platform and DNA extraction according to the manufacturers protocol. The DNA concentration of extracts was checked at random using a High Sensitivity dsDNA Kit on a Qubit fluorometer (Thermo Fisher Scientific). In total, we compiled 642 DNA extracts of 261 species.

#### 1.1. Sanger sequencing

For the design of Sanger sequencing primers, regions of low interspecific variability upstream and downstream of the selected priming sites were chosen. The two newly designed Ave-seq primer pairs showed minimum secondary structures, included ambiguous bases for maximum coverage across bird species, and had melting temperatures between 50 and 55 °C using the OligoAnalyzer^TM^ Tool (Integrated DNA Technologies; Table 1).

**Table 1:**
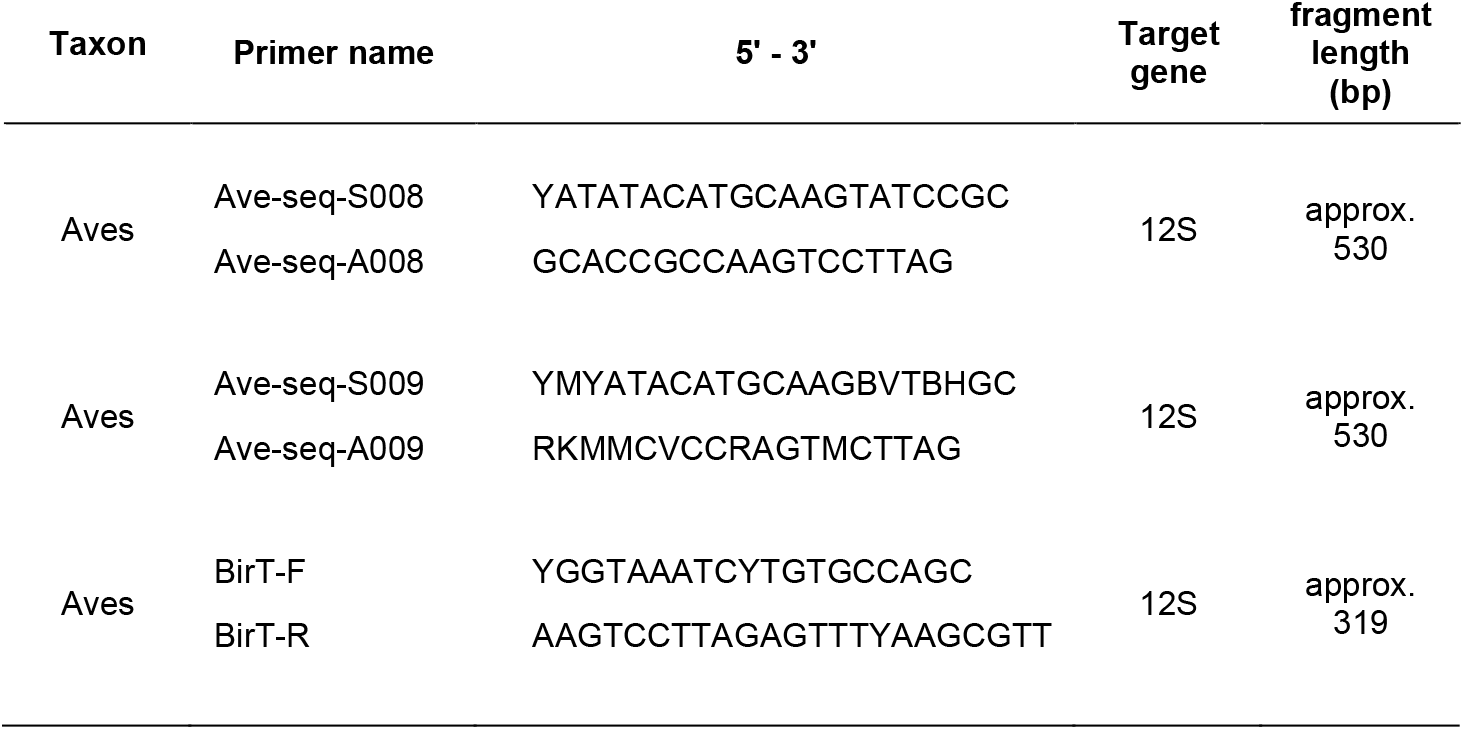
The primers developed for Sanger Sequencing (Ave-seq) and 12S eDNA metabarcoding (BirT)

All DNA extracts generated from bird tissues were initially amplified with the Ave-seq-S008 / Ave-seq-A008 primer pair. The 15 µl reactions each contained 2 × reaction mix from the Multiplex PCR Kit (Qiagen), 5 µg Bovine Serum Albumin, 0.5 µM of each primer, 3 µl of DNA extract, and molecular grade water. Optimized thermocycling conditions were 15 min at 95 °C, 35 cycles of 30 s at 94 °C, 60 s at 52 °C, 1 min at 72 °C, and 10 min at 72 °C once. Amplification success was checked on 1% agarose gels. If no amplification success was observed, PCR was repeated under the same conditions with primers Ave-seq-S009 / Ave-seq-A009. Subsequently, all PCR products, independent of amplification success, were cleaned using SPRIselect (Beckman Coulter) with a sample to volume ratio of 1.5 × and Sanger-sequenced at the AAC Genomics Facility (University of Guelph). The generated sequences were manually curated to ensure base call quality and remove ambiguities in Geneious Prime® before uploading them to BOLD (Project BTBIR; SI2). For further analysis, their names were adjusted to the IOC conventions and changes in the coverage of the reference sequence database were plotted.

Ultimately, a custom reference database (SI3) was compiled which contained all newly generated 12S reference sequences and the previously available 12S bird sequences both of which were required to cover 100% of the amplified fragment and could be aligned to *B. canadensis*. Of the 5,021 sequences contained in this database, 473 were associated with more than one taxonomic entity on the subspecies or species level. These ambiguities were noted for the correct interpretation of any field-derived results.

### eDNA samples: field collection

Water samples were taken every other day around noon at the Mountsberg conservation area (43.455094, −80.046249) in Southern Ontario between the 6^th^ and 14^th^ November 2019. Due to cold weather conditions, the lake was partially frozen, but waterfowl were still present on all days. To avoid ice cover, samples were taken close to the outflow integrated into the artificial dam. Water was filtered using three different filter types: Nature Metrics filters with 47 mm diameter and GF (5 µm) plus PES (0.8 µm) membranes (hereafter “NM filters”) were operated with 60 ml syringes. Self-preserving Smith Root filters with 47 mm diameter and a PES membrane of 1.2 µm or 5 µm pore size (hereafter “SM filters”), which were processed with an eDNA Sampler Backpack (Smith Root) at standard settings. First, 15 l of water were taken with a bucket from the lake. Three liters of bucket water were filtered across each filter type. Afterwards, the bucket was emptied, rinsed with 5% bleach and deionized water, before refilling it with lake water and repeating the filtration procedure another two times to a total of 9 filters (3 per filter type). In case the 3 l could not be reached due to elevated levels of turbidity, the filtered water volume was noted (SI4). After every second sampling day, one liter of deionized water was filtered across one filter of each type in the field after processing the lake water. Upon returning to the University of Guelph, the NM filters were immediately frozen at −20 °C until further processing. The SR filters were left in their packaging at room temperature over night to enable drying of the filter membrane, before also storing them at −20 °C. DNA-free gloves were worn during filtration and changed between samples. The backpack sampler was cleaned and calibrated according to the manual before each sampling day and all multi-use equipment was cleaned for 10 min with 5% bleach and rinsed with deionized water.

### eDNA samples: laboratory processing

After thawing, 600 µl of lysis buffer (TES (0.1 M TRIS, 10 mM EDTA, 2% sodium dodecyl sulphate; pH 8) and Proteinase K (Qiagen, 20 mg/ml) in a 20:1 ratio) were added to the NM filters. SM filters were placed in 2 ml reaction tubes together with 200 µl of lysis buffer. All filters were incubated over night at 56°C. The lysis buffer from the NM filters was transferred to a 2 ml reaction tube by pushing air into the filter capsule with a syringe, while the buffer was centrifuged out of the SM filters into a 2 ml reaction tube for 10 min at 20,000 × g using a plastic inlet with perforated bottom (Thalinger, Kirschner, et al., 2021). From this step on, we followed the standard DNeasy Blood and Tissue protocol for all samples with two modifications: i) total lysis buffer volumes were noted for all samples and equal volumes of buffer AL and ethanol were added. In case the total volume of the mixture exceeded 900 µl, consecutive binding steps were used to process the entire lysate volume. ii) Two elution steps with 75 µl buffer AE each were carried out. For each filter type, one extraction control containing only lysis buffer was processed along the other samples. Next, the total DNA content of all extracts was quantified using qubit (SI4).

For bird eDNA metabarcoding, all extracts (including field and extraction controls) were subjected to two consecutive PCRs using a fusion primer approach (Elbrecht & Steinke, 2018) and optimized PCR conditions (SI5). PCRs were carried out in 96-well plates and each row contained one randomly positioned PCR negative control. The first PCR was carried out with the BirT primers (Table 1): each 20 µl reaction contained 10 µl of 2 × Multiplex Master Mix, 1 µl of each primer (10 µM), 4 µl DNA extract, and molecular grade water. Thermocycling conditions were 10 min of initial denaturation at 95°C, followed by 25 cycles of 95°C for 30 s, 60°C for 30 s, and 72°C for 1 min, and final extension at 72°C for 10 min. All PCR products were cleaned up using a SPRIselect (Beckman Coulter) with a 1.5 × ratio between PCR product and SPRI beads, washing with 70% ethanol, and elution in 10 µl molecular grade water. Afterwards, the second PCR was carried out in 20 µl reactions with 10 µl of 2 × Multiplex Master Mix, 1 µl of the forward and reverse fusion primers (10 µM; including Illumina sequencing adapters; SI6), 2 µl of the cleaned-up PCR product, and molecular grade water. The thermocycling conditions were the same as for the first PCR, except for an annealing temperature of 50 °C. Successful amplification was confirmed by visualizing amplicons on 1.5% agarose gels.

SequalPrep Normalization Plates (Thermo Fisher Scientific) were used to purify 15 µl of each PCR product according to the manufacturer’s instructions (Harris et al., 2010). After normalization, 10 µl from each well were pooled. Consecutively, left-sided and right-sided size selection using SPRIselect were carried out with ratios of 0.7 and 0.6, respectively. The final library was checked on an 1.5% agarose gel and its average concentration (5 measurements) determined to be 5.09 ng/µl via qubit. Sequencing was carried out at the McMaster University, Hamilton using a NovaSeq SP flow cell with the 500 cycle kit (Illumina) and 5% PhiX spike in. The samples were sequenced together with another bird eDNA library aiming at a sequencing depth of at least 800,000 reads per sample.

### eDNA samples: sequence processing

Raw sequences were processed with JAMP (https://github.com/VascoElbrecht/JAMP) and data from the two lanes of the flow cell were kept separate. During demultiplexing, only sequences with perfectly matching tags were considered for further processing. Demultiplexed sequencing results were uploaded to the Sequence Read Archive (SRA) and will become available upon acceptance. Usearch (Edgar, 2010) was used to merge forward and reverse reads requiring at least a 75% match. Primer sequences were removed with Cutadapt (default settings; Martin, 2011) and afterwards, only fragments with a length of 265 bp ± 30 bp were considered for further processing. To remove sequences with poor quality, an expected error value of 1 was applied via Usearch (Edgar & Flyvbjerg, 2015). All amplicon sequence variants (ASV) with less than 3 reads per sample and lane or less than 0.001% relative abundance in at least one sample were removed. The remaining ASVs were mapped twice against a custom reference database: once against all bird sequences (SI3) and non-targets (SI1) with a minimum required match of 85% and once against the custom bird sequence database alone (SI3) requiring a 100% match to enable a direct assignment at species level in as many cases as possible (Elbrecht & Leese, 2017b). The first mapping was carried out to examine the likelihood of non-target amplification, the second to taxonomically assign the obtained reads to bird species. This was followed by a manual plausibility check. Subsequently, genus-level information was used if the taxonomic resolution in the reference database was ambiguous, and more than one species occurred in the study area.

Prior to analysis, reads in filtration, extraction and PCR control were examined. Since all control types contained low levels of contamination with target DNA (median read number in control samples was 0.88% of the median read number in field samples), a strict contamination correction regime was applied: for every bird species the highest number of reads detected in a filtration, extraction, and PCR control, respectively, were added. From each field sample containing reads of the species in question, twice this read number was subtracted. Since the sequence processing and bioinformatic analysis was carried out separately for the two lanes, this correction was also applied separately.

### eDNA samples: data analysis

For a comparison between the eDNA-based results and data collected by birdwatchers, seven bird watching checklists submitted to eBird (https://eBird.org) over the last week of October and the first two weeks of November 2019 for the locations “Mountsberg Wellington” and “Mountsberg Hamilton” were downloaded. All calculations and visualizations of the eDNA results were made in R Version 4.2.2 (R Core Team, 2022) using the packages “ggplot2” (H. Wickham, 2016), “dplyr” (Hadley Wickham, François, Henry, & Müller, 2019), “ggpubr” (Kassambara, 2019), “ggsignif” (Ahlmann-Eltze & Patil, 2021), and “randomcoloR” (Ammar, 2019). Poisson regressions with a log-link function were used to test for significant differences in read numbers and the number of detected taxa between filter types and a Kruskal-Wallis ANOVA followed by Wilcoxon-signed-rank tests were employed to detect pairwise differences in filtered water volume between filter types.

## Results

### BirT Primers

The 12S fragment amplified by the BirT-F and BirT-R primers (Table 1) is approximately 320 bp long (including primers) and the fragment including all tags and sequencing adaptors after PCR 2 has a length of approx. 470 bp. The BirT-F and BirT-R primers are 19 and 23 bp long, have two and one degenerate bases, and average melting temperatures of 57.4 °C and 57.2 °C, respectively. The BirT-F primer has 1-4 mismatches to non-target groups, all of which are located in the middle of the priming site. The BirT-R primer has at least three mismatches, and at least one of them is located in the first three bases at the 3’ end (Fig. 2). These mismatches in combination with an optimized annealing temperature of 60 °C in the first PCR minimize amplification of non-target taxa. Other 12S bird metabarcoding primers encompass one or two regions of high sequence variability, while the BirT primer fragments encompasses four such regions, thus improving the chance to identify individual species at comparably short fragment length (Fig. 1). Albeit the exact location and primer sequences differ, the BirT-F and the MiBird-U-F primer (Ushio et al., 2018) are placed in the same region of 12S. Additionally, the BirT-R primer has some overlap with the Aves_12Sc (EPP et al., 2012) and Aves01-R (Taberlet et al., 2018) primers (Fig. 1).

### Sequence reference database

For the 10933 non-extinct bird species listed in the IOC World Bird List, 4,443 12S sequences were available, which spanned the entire BirT fragment. For 8,790 of species (80.3%) a reference sequence was not available (Fig. 3). For 6.8% (n = 744) and 6.6% (n = 726) of the species one or two reference sequences were available, respectively, and 3-18 sequences were attributed to the remaining species. Most bird species belong to the order of Passeriformes for which only 12,9% (n = 851) of species were associated with a reference sequence. For the majority of orders, the percentage of species with available reference sequences was between 20 and 50% (Fig. 3). Sanger sequencing resulted in the generation of 594 novel 12S sequences belonging to 254 species (BOLD Project BTBIR; SI2). For 578 sequences both DNA strands were available across the length of the entire fragment and thus added to the final reference database which in the end contained 5021 sequences of 2143 species (SI3). Charadriiformes and Anseriformes were the orders for which most new sequences were generated and Charadriiformes and Gaviiformes the orders for which the new sequences most improved the percentage of species with available reference sequences (Fig. 3).

**Figure 3:**
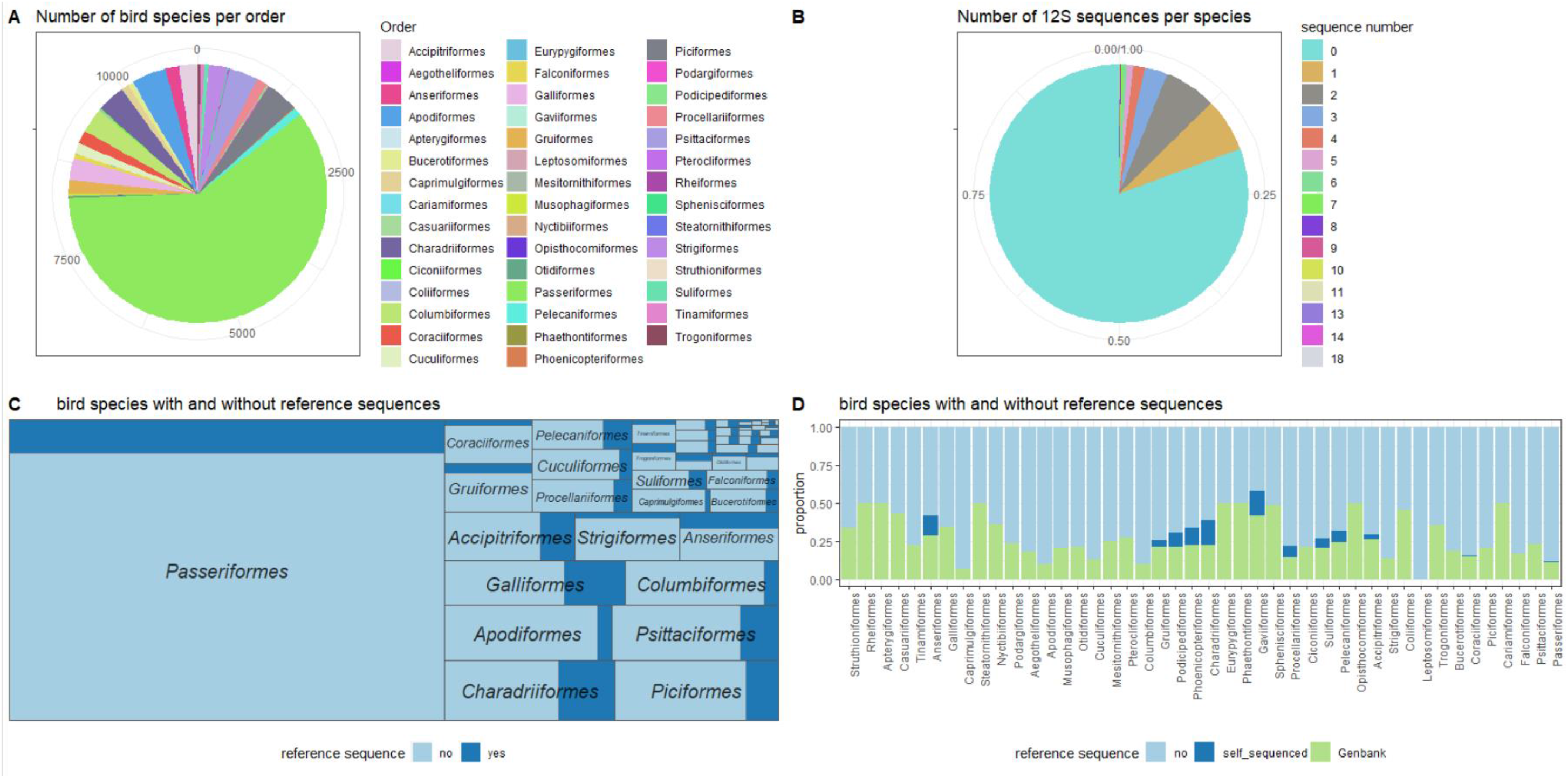
Panel A summarizes the number of extant bird species by order while panel B provides an overview of the number of the publicly available 12S sequences (as of Jan 2023) which cover the entire fragment amplified by the BirT primer pair. Panel C displays how these sequences are associated with each order and displays the ratio of species for which a public reference sequence was available. Finally, Panel D displays the changes in the publicly available 12S bird sequences that were made by the Sanger sequencing efforts in this publication.

### eDNA samples

After demultiplexing, 58,639,599 paired-end reads of the NovaSeq SP run were associated with the eDNA samples and controls relevant to this publication. After merging, removal of the primer sequences, length selection, quality filtering, and denoising 32,345,885 reads (sum of the two lanes) were assigned with at least an 85% match to either a bird or a non-target species, only 253 reads were identified as reptile sequences. The 27,056,563 reads which were assigned with a 100% match to the bird reference sequence database, were used for all subsequent analyses. Accordingly, the average sequencing depth of field-collected eDNA samples after correcting for contaminations was 613,972 ± 340,088 SD reads. A taxonomic assignment to a species with high plausibility of occurring in the area was possible for 95% of these reads and 24 taxa (22 at species-level, two at genus level) were detected in 42 of the 48 eDNA samples. An effect of filter type could not be detected, neither for read number per sample nor target species per sample (mean: 5.5 ± 2.3 SD) (Fig. 4). On average, 2.2 l ± 0.96 l SD of water were filtered, with the Smith-Root 1.2 µm filters allowing for significantly less filtered water volume per sample than the other two filter types (Fig.4).

**Figure 4:**
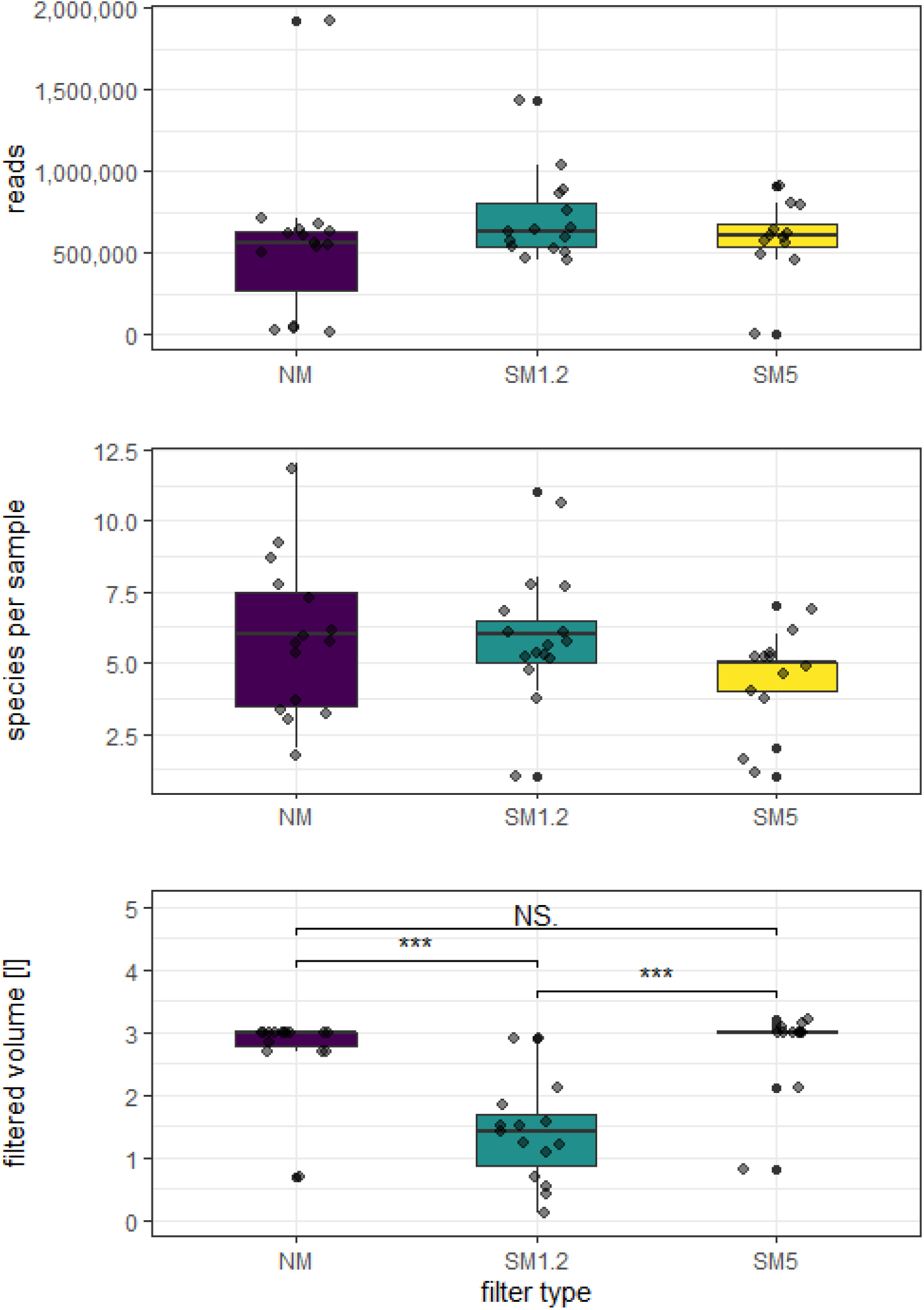
The top panel show how reads per taxon are distributed between filter types. The Middle panel displays the number of detected species per sample and filter type (data in both panels based on 100% matches with the reference database; SI3). The bottom panel displays the filtered water volume per sample and filter type. Based on Poisson regressions with a log-link function, no significant differences between filter types were detected for read and species numbers per sample. However, a Kruskal-Wallis Anova followed by Wilcoxon-signed-rank tests detected a significantly lower volume of water filtered for the Smith-Root 1.2 µm filters (SM1.2) in comparison to the Nature metrics filters (NM) and the Smith-Root 5 µm filters (SM5). *** codes for p<0.001 and N.S. for “not significant”.

In most samples, the majority of reads were assigned to *Branta canadensis* and *Anas* spp. with the former dominating samples from 8^th^, 10th and 12^th^ Nov, and the latter in samples obtained on 6^th^ and 14^th^ November (Fig. 5, Fig. 6). Additionally, *Mareca americana* and *Fulica americana* were detected in 32 and 23 samples, respectively. On most occasions, the detected species communities per sample were highly similar between samples obtained from the same bucket of water, with the most notable exception being two Smith-Root 5µm filters obtained on the final sampling date in which *Coturnicops noveboracensis* and *Porzana carolina* contributed the majority of reads (Fig. 5). Fifty-eight percent of the highly plausible detected taxa were also recorded in the visual checklists uploaded to eBird, with Anseriformes being the most frequently represented order (Fig. 6). Only two of the species, *Turdus migratorius* and *Catharus guttatus,* did not have a close association with aquatic habitats. Of the 34 bird species listed in the visual checklists but not detected in the eDNA samples, 70.5% were not closely associated with aquatic habitats and 53% belonged to the order Passeriformes.

**Fig. 5:**
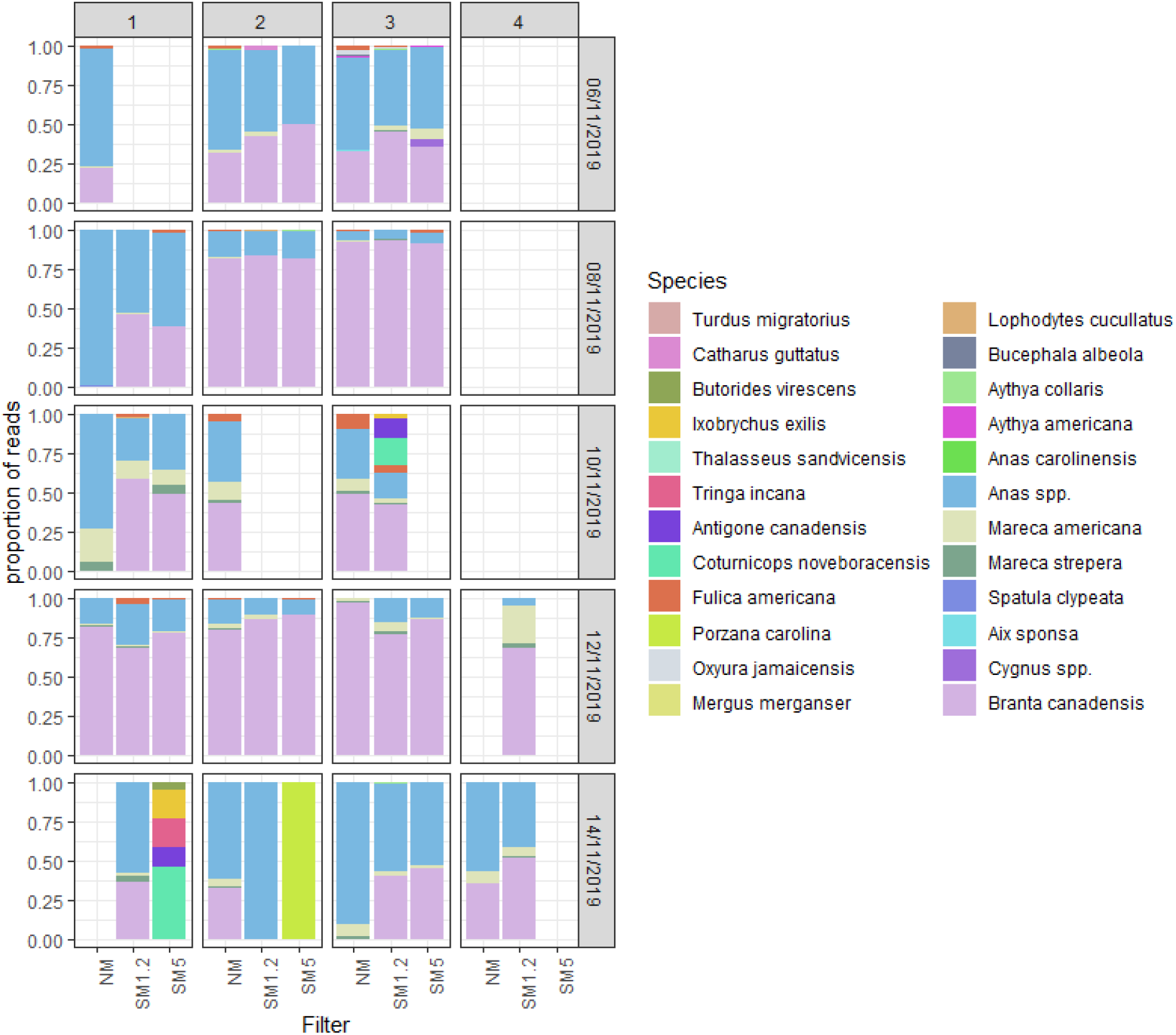
Bar chart displaying the bird taxa with a 100% match to the reference database (SI3). Each bar denotes the proportion of reads per sample and species. Columns 1 to 4 code for the bucket number from which one Nature metrics filter (NM), one Smith-Root 1.2 µm filters (SM1.2) and one Smith-Root 5 µm filter (SM5) were processed in most cases, respectively. Rows denote distinct sampling dates from 6^th^ November 2019 to 14^th^ November 2019.

**Fig. 6:**
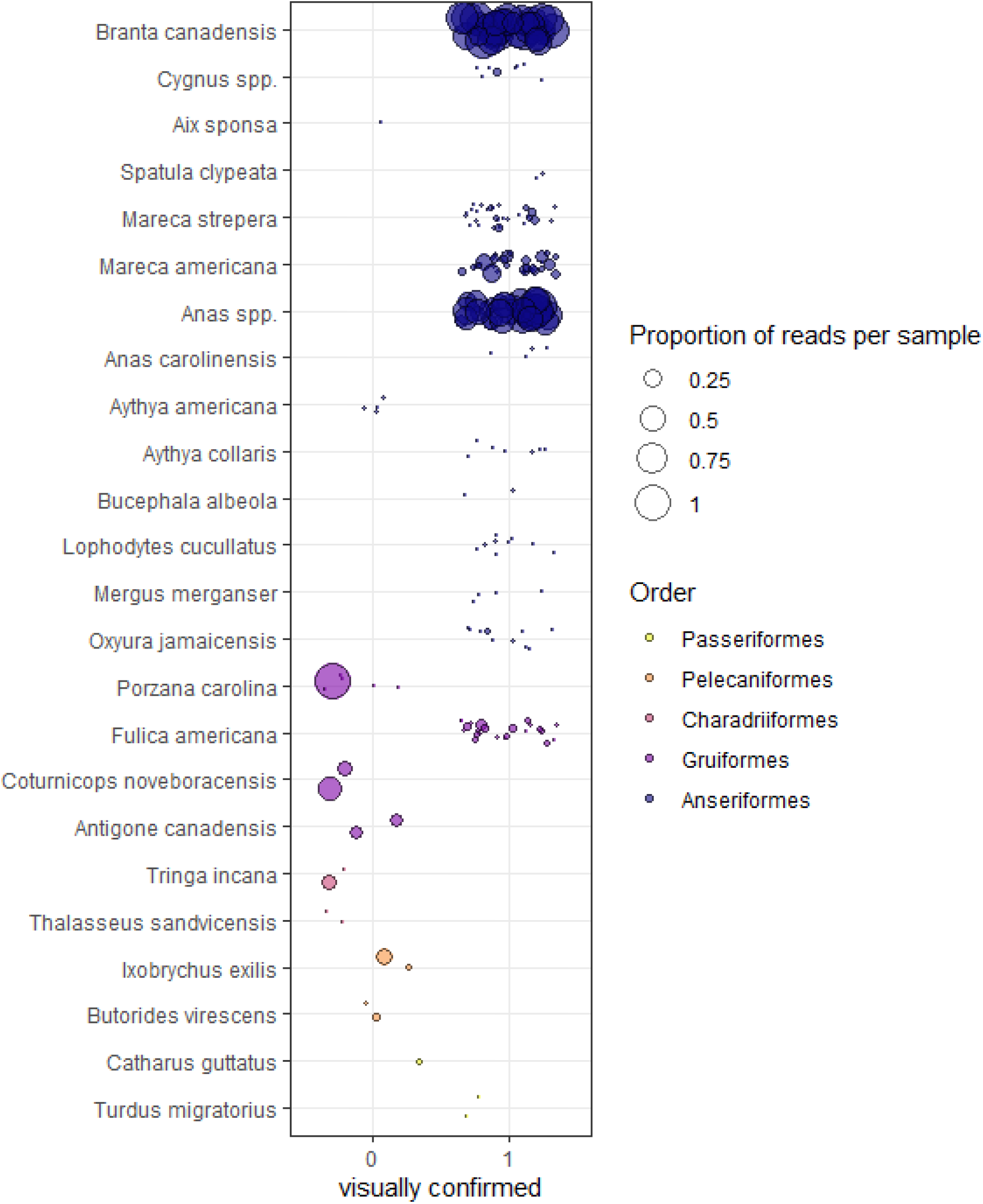
The proportion of reads per sample assigned to each of the detected taxa. Circle size represents read proportions, circle color the order for each taxon. Additionally, detections are divided based on whether a visual confirmation was contained in checklists uploaded to eBird during the sampling period with “0” coding for no confirmation and “1” coding for confirmation. Of the detected taxa, only *Catharus guttatus* and *Turdus migratorius* are not closely associated with water.

## Discussion

The results obtained from field-collected eDNA samples confirm the suitability of the BirT primer pair for avian eDNA metabarcoding with no detectable amplification of non-targets such as mammals, fishes, or amphibians. The length of the amplified fragment (approx. 320 bp after first PCR, approx. 470 bp after second PCR) allows sequencing with kits designed for at least 500 cycles on Illumina instruments while still providing sufficient overlap between forward and reverse reads (Deiner et al., 2017). Additionally, the four high variability regions within the amplified region maximize the chances of species-specific matches to reference barcodes. Thus, the primer pair strikes the ideal balance between a fragment length short enough for the detection of potentially degraded eDNA, suitability for high throughput sequencing, minimal non-target amplification, and the generation of datasets with sufficient taxonomic depth to enable the monitoring of birds via eDNA on a species-specific level.

For the application of the BirT primers with eDNA from filtered water samples, the mismatches hindering the amplification of fishes are of particular importance, since most primer pairs targeting vertebrates were originally developed for the detection of fish species (Zhang et al., 2020; Mariani et al., 2021). For the growing number of studies investigating bird eDNA from filtered air samples, mismatches at the priming site avoiding the amplification of human DNA are of particular importance (Clare et al., 2021; Johnson et al., 2023; Lynggaard et al., 2023). The non-amplification of human DNA is also key for any studies employing a community science approach, in which measures to avoid the contamination of samples such as frequent change of gloves, are not always feasible.

Targeting fragments of the mitochondrial 12S gene for the detection of vertebrates in environmental samples has frequently provided superior taxonomic resolution and taxon-specific amplification in comparison with other potential target genes such as the COI (Deagle et al., 2014; Collins et al., 2019). Unfortunately, the incompleteness of the sequence reference database still has negative effects on the taxonomic resolution, since originally, vertebrate barcoding efforts have been focused on COI and CytB genes (Lavinia, Kerr, Tubaro, Hebert, & Lijtmaer, 2016). Our analysis showed, that for 80.3% of all extant bird species, a 12S sequence is not available. Additionally, changes in bird taxonomy and differences in accepted nomenclature complicate the assembly of reference databases (e.g. Lepage, Vaidya, & Guralnick, 2014; Donsker & Rasmussen, 2022). Our own efforts to increase the number of available 12S reference sequences resulted in the generation of 594 new DNA barcodes representing 254 species. Due to the location of the field sampling site in southern Ontario and the processing of water samples, we focused on the generation of new barcodes for bird species occurring in Canada associated with aquatic habitats. The final curated reference sequence dataset (SI3) is not only the largest bird 12S reference database to date, but it also accounts for taxonomic ambiguities (i.e. cases in which species can not be resolved to species level). Ultimately, only a systematic international effort, analog to the original COI barcoding efforts (Hebert et al., 2003) but targeting 12S or entire mitogenomes could help to close the substantial gaps in the reference database and facilitate future taxonomic assignments independent of geographic sampling location and habitat type.

Recently, the use of eDNA data has gained attention for applications going beyond the detection of individual species (Yates et al., 2023). For example, Elbrecht et al. (2018) investigated the presence of distinct haplotypes for arthropod bulk samples collected in Finland, and Dugal et al. (2021) were able to haplotype whale sharks (*Rhincodon typus*) from eDNA samples. In the present study, taxonomic mapping was not employed beyond the species level. However, 302 entries in the sequence reference database are resolved to the sub-species level (Passeriformes and Charadriiformes with 66 and 50 sequences, respectively). Additionally, for 65.2% of the species with 12S DNA barcodes, more than one reference sequence is available. This opens the door for future efforts to test the BirT fragment for its suitability to detect subspecies and haplotypes; efforts which usually only rely on sequence variations detected in the newly generated metabarcoding datasets (Elbrecht et al., 2018).

Our analysis of field-collected eDNA samples with the BirT primers is a proof of concept for the applicability of the primer pair and the reference database for bird eDNA metabarcoding. The strict 100% threshold for taxonomic assignment at the species level (c.p. Lynggaard et al., 2023) was deliberately chosen to minimize the influence of sequencing errors and contamination, thus potentially forgoing the taxonomic assignment of some low read number ASVs for which a perfect match with the reference database was not possible. The majority of all taxonomically matched reads (95%) passed the manual plausibility check, i.e. the detected species has been known to occur in the study region. For the remaining 5%, the detection was either unlikely based on data available on eBird, or a closely related species was known to occur in the area allowing for a taxonomic assignment only to genus level.

Albeit the filtered water volume significantly varied between samples, an effect of filter type (i.e. NM filters, SM 1.2µm and SM 5µm filters), and the associated filtration method (syringe vs. pump) on read number per sample and number of detected species per sample could not be observed. For the detection of eDNA, the selection of a suitable pore size depends on features of the sampled habitat (e.g. prevailing levels of turbidity) and the state in which the target DNA is present in the environment (Jo et al., 2022; Mauvisseau et al., 2022). Previously, pore sizes between 0.2 and 5µm have been found suitable for the efficient detection of vertebrate eDNA from water samples (Tsuji, Takahara, Doi, Shibata, & Yamanaka, 2019); particularly in situations with limited turbidity. Albeit sampling was carried out late in the year with large parts of the sampling site already covered by ice, the water samples exhibited notable levels of turbidity and algae, which led to filtration of less than 2.5 l for 33.3% of the samples. Nevertheless, we did not find a negative correlation between filtered water volume and read numbers or the number of detected species. This shows that all selected filters, filtration methods, and water volumes were suitable to detect bird species which are closely associated with aquatic habitats utilizing their eDNA.

The species and genera which were detected with eDNA and confirmed by visual observations during the field sampling period, all occurred in more than one sample. *B. canadensis* and *Anas* spp. Were most frequently detected, had the highest percentage of reads associated with them, and were most abundant in visual observations. Additionally, the presence of these species on the ice or in the water was noted on all sampling dates. Generally, the detected species match the bird species observed during fall migration in this part of Ontario when compared with visual checklists uploaded to eBird. Due to the limited availability of visual bird observations collected by citizen scientists – especially for remote locations and when only a small time window is considered and during migration (Kamp et al., 2016) – the discrepancy between the eDNA-based and observation-based datasets is not surprising.

## Conclusion

This pilot study confirms the suitability of the BirT primer pair for bird eDNA metabarcoding with optimized fragment length, good taxonomic resolution, and mismatches to non-target groups. Although there are still substantial gaps in the 12S reference sequence database, especially for less common species which primarily occur outside of temperate regions, the analysis of bird eDNA from water samples collected in southern Ontario confirmed the applicability of the approach for all tested filter/filtration combinations. Bird detections based on 100% matches with the reference sequence database were highly plausible and were confirmed by visual bird observations uploaded to eBird during the sampling period. The majority of the detected bird species from water samples is closely associated with aquatic habitats (Ushio et al., 2018), whilst the analysis of eDNA from air samples primarily results in the detection of bird species associated with terrestrial habitats (Littlefair et al., 2023; Lynggaard et al., 2023). This indicates that in the future, a combined analysis of water and air samples might be ideal to provide a complete picture of local bird diversity using eDNA.

## Acknowledgments

We thank the Mountsberg conservation authority for the permission to collect eDNA samples. Additionally, we are very grateful to Doug Touzer for providing taxonomic expertise on Canadian Bird species and the team at the Royal Ontario Museum (M. Peck and S. Claramut), who allowed us to subsample their bird tissue collection. A big “thank you” also goes to C. Hempel, N. Houde, E. Vilacoba and P Eder for their help with tissue and eDNA sampling.

## Data availability

The raw metabarcoding data will be available at GenBank SRA upon publication, the generated 12S barcodes are available at BOLD (Project BTBIR) and in SI2. All other datasets are available as Supporting Information files.

## Author contribution statement

BT conceived the original idea, which was developed further together with DS who secured the necessary funding. BT was responsible for tissue collection and eDNA sampling, the analysis of the available 12S bird sequences and primer design. Under the supervision of BT, RE, MC, and KC contributed significantly to the laboratory processing of all samples. BT analyzed the obtained datasets and wrote a first draft of the manuscript to which all co-authors contributed before final approval for publication.

## References

Ahlmann-Eltze, C., & Patil, I. (2021). ggsignif: R Package for Displaying Significance Brackets for “ggplot2.” PsyArxiv. doi:10.31234/osf.io/7awm6

Aide, T. M., Corrada-Bravo, C., Campos-Cerqueira, M., Milan, C., Vega, G., & Alvarez, R. (2013). Real-time bioacoustics monitoring and automated species identification. PeerJ, 2013(1), e103. doi:10.7717/peerj.103

Ammar, R. (2019). randomcoloR: Generate Attractive Random Colors. Retrieved from https://cran.r-project.org/package=randomcoloR

Bista, I., Carvalho, G. R., Tang, M., Walsh, K., Zhou, X., Hajibabaei, M., … Creer, S. (2018). Performance of amplicon and shotgun sequencing for accurate biomass estimation in invertebrate community samples. Molecular Ecology Resources, 18(5), 1020–1034. doi:10.1111/1755-0998.12888

Bohmann, K., & Lynggaard, C. (2023, February 1). Transforming terrestrial biodiversity surveys using airborne eDNA. Trends in Ecology and Evolution. Elsevier Ltd. doi:10.1016/j.tree.2022.11.006

Bruce, K., Blackman, R., Bourlat, S. J., Hellström, A. M., Bakker, J., Bista, I., … Deiner, K. (2021). A practical guide to DNA-based methods for biodiversity assessment. A practical guide to DNA-based methods for biodiversity assessment. Pensoft Publishers. doi:10.3897/ab.e68634

Clare, E. L., Economou, C. K., Faulkes, C. G., Gilbert, J. D., Bennett, F., Drinkwater, R., & Littlefair, J. E. (2021). eDNAir: Proof of concept that animal DNA can be collected from air sampling. PeerJ, 9. doi:10.7717/peerj.11030

Collins, R. A., Bakker, J., Wangensteen, O. S., Soto, A. Z., Corrigan, L., Sims, D. W., … Mariani, S. (2019). Non-specific amplification compromises environmental DNA metabarcoding with COI. Methods in Ecology and Evolution, 10(11), 1985–2001. doi:10.1111/2041-210X.13276

Deagle, B. E., Jarman, S. N., Coissac, E., Pompanon, F., & Taberlet, P. (2014). DNA metabarcoding and the cytochrome c oxidase subunit I marker: Not a perfect match. Biology Letters, 10(9). doi:10.1098/rsbl.2014.0562

Deiner, K., Bik, H. M., Mächler, E., Seymour, M., Lacoursière-Roussel, A., Altermatt, F., … Bernatchez, L. (2017). Environmental DNA metabarcoding: Transforming how we survey animal and plant communities. Molecular Ecology, 26(21), 5872–5895. doi:10.1111/mec.14350

Donsker, G. F. D., & Rasmussen, P. (2022). IOC World Bird List (v12.2). Retrieved from https://doi.org/10.14344/IOC.ML.12.2.

Dugal, L., Thomas, L., Jensen, M. R., Sigsgaard, E. E., Simpson, T., Jarman, S., … Meekan, M. (2021). Individual haplotyping of whale sharks from seawater environmental DNA. Molecular Ecology Resources, 1755–0998.13451. doi:10.1111/1755-0998.13451

Edgar, R. C. (2010). Search and clustering orders of magnitude faster than BLAST. Bioinformatics, 26(19), 2460–2461. doi:10.1093/bioinformatics/btq461

Edgar, R. C., & Flyvbjerg, H. (2015). Error filtering, pair assembly and error correction for next-generation sequencing reads. Bioinformatics, 31(21), 3476–3482. doi:10.1093/bioinformatics/btv401

Elbrecht, V., Braukmann, T. W. A., Ivanova, N. V., Prosser, S. W. J., Hajibabaei, M., Wright, M., … Steinke, D. (2019). Validation of COI metabarcoding primers for terrestrial arthropods. PeerJ, 2019(10), e7745. doi:10.7717/peerj.7745

Elbrecht, V., & Leese, F. (2015). Can DNA-Based Ecosystem Assessments Quantify Species Abundance? Testing Primer Bias and Biomass—Sequence Relationships with an Innovative Metabarcoding Protocol. PLOS ONE, 10(7), e0130324. doi:10.1371/journal.pone.0130324

Elbrecht, V., & Leese, F. (2017a). PrimerMiner: an R package for development and in silico validation of DNA metabarcoding primers. Methods in Ecology and Evolution, 8(5), 622–626. doi:10.1111/2041-210X.12687

Elbrecht, V., & Leese, F. (2017b). Validation and Development of COI Metabarcoding Primers for Freshwater Macroinvertebrate Bioassessment. Frontiers in Environmental Science, 5(APR), 11. doi:10.3389/fenvs.2017.00011

Elbrecht, V., & Steinke, D. (2018). Scaling up DNA metabarcoding for freshwater macrozoobenthos monitoring. Freshwater Biology, 64(2), fwb.13220. doi:10.1111/fwb.13220

Elbrecht, V., Vamos, E. E., Steinke, D., & Leese, F. (2018). Estimating intraspecific genetic diversity from community DNA metabarcoding data. PeerJ, 2018(4). doi:10.7717/peerj.4644

Epp, L. S., Boessenkool, S., Bellemain, E. P., Haile, J., Esposito, A., Riaz, T., … Brochmann, C. (2012). New environmental metabarcodes for analysing soil DNA: potential for studying past and present ecosystems. Molecular Ecology, 21(8), 1821–1833. doi:10.1111/j.1365-294X.2012.05537.x

Ficetola, G. F., & Taberlet, P. (2023). Towards exhaustive community ecology via DNA metabarcoding. Molecular Ecology. doi:10.1111/mec.16881

Harris, J. K., Sahl, J. W., Castoe, T. A., Wagner, B. D., Pollock, D. D., & Spear, J. R. (2010). Comparison of normalization methods for construction of large, multiplex amplicon pools for next-generation sequencing. Applied and Environmental Microbiology, 76(12), 3863–3868. doi:10.1128/AEM.02585-09

Hebert, P. D. N., Cywinska, A., Ball, S. L., & DeWaard, J. R. (2003). Biological identifications through DNA barcodes. Proceedings of the Royal Society B: Biological Sciences, 270(1512), 313–321. doi:10.1098/rspb.2002.2218

Holman, L. E., de Bruyn, M., Creer, S., Carvalho, G., Robidart, J., & Rius, M. (2021). Animals, protists and bacteria share marine biogeographic patterns. Nature Ecology and Evolution, 5(6), 738–746. doi:10.1038/s41559-021-01439-7

Jackman, J. M., Benvenuto, C., Coscia, I., Oliveira Carvalho, C., Ready, J. S., Boubli, J. P., … Guimarães Sales, N. (2021). eDNA in a bottleneck: Obstacles to fish metabarcoding studies in megadiverse freshwater systems. Environmental DNA, 3(4), 837–849. doi:10.1002/edn3.191

Jensen, M. R., Høgslund, S., Knudsen, S. W., Nielsen, J., Møller, P. R., Rysgaard, S., & Thomsen, P. F. (2023). Distinct latitudinal community patterns of Arctic marine vertebrates along the East Greenlandic coast detected by environmental <scp>DNA</scp>. Diversity and Distributions, 29(2), 316–334. doi:10.1111/ddi.13665

Jo, T., Takao, K., & Minamoto, T. (2022, March 1). Linking the state of environmental DNA to its application for biomonitoring and stock assessment: Targeting mitochondrial/nuclear genes, and different DNA fragment lengths and particle sizes. Environmental DNA. John Wiley and Sons Inc. doi:10.1002/edn3.253

Johnson, M. D., Barnes, M. A., Garrett, N. R., & Clare, E. L. (2023). Answers blowing in the wind: Detection of birds, mammals, and amphibians with airborne environmental DNA in a natural environment over a yearlong survey. Environmental DNA, 5(2), 375–387. doi:10.1002/edn3.388

Kamp, J., Oppel, S., Heldbjerg, H., Nyegaard, T., & Donald, P. F. (2016). Unstructured citizen science data fail to detect long-term population declines of common birds in Denmark. Diversity and Distributions, 22(10), 1024–1035. doi:10.1111/ddi.12463

Kassambara, A. (2019). ggpubr: “ggplot2” Based Publication Ready Plots. Retrieved from https://cran.r-project.org/package=ggpubr

King, R. A., Read, D. S., Traugott, M., & Symondson, W. O. C. (2008). Molecular analysis of predation: A review of best practice for DNA-based approaches. Molecular Ecology, 17(4), 947–963. doi:10.1111/j.1365-294X.2007.03613.x

Klure, D. M., Greenhalgh, R., & Dearing, M. D. (2022). Addressing nontarget amplification in DNA metabarcoding studies of arthropod-feeding rodents. Mammal Research, 67(4), 499–509. doi:10.1007/s13364-022-00646-2

Lavinia, P. D., Kerr, K. C. R., Tubaro, P. L., Hebert, P. D. N., & Lijtmaer, D. A. (2016). Calibrating the molecular clock beyond cytochrome *b* : assessing the evolutionary rate of COI in birds. Journal of Avian Biology, 47(1), 84–91. doi:10.1111/jav.00766

Lepage, D., Vaidya, G., & Guralnick, R. (2014). Avibase - A database system for managing and organizing taxonomic concepts. ZooKeys, 420(420), 117–135. doi:10.3897/zookeys.420.7089

Levesque-Beaudin, V., Steinke, D., Böcker, M., & Thalinger, B. (2023). Unravelling bird nest arthropod community structure using metabarcoding. BioRxiv, 2023.03.09.531929. doi:10.1101/2023.03.09.531929

Li, F., Qin, S., Wang, Z., Zhang, Y., & Yang, Z. (2023). Environmental DNA metabarcoding reveals the impact of different land use on multitrophic biodiversity in riverine systems. Science of the Total Environment, 855, 158958. doi:10.1016/j.scitotenv.2022.158958

Littlefair, J. E., Allerton, J. J., Brown, A. S., Butterfield, D. M., Robins, C., Economou, C. K., … Clare, E. L. (2023). Air-quality networks collect environmental DNA with the potential to measure biodiversity at continental scales. Current Biology, 33(11), R426–R428. doi:10.1016/j.cub.2023.04.036

Lynggaard, C., Frøslev, T. G., Johnson, M. S., Olsen, M. T., & Bohmann, K. (2023). Airborne environmental <scp>DNA</scp> captures terrestrial vertebrate diversity in nature. Molecular Ecology Resources. doi:10.1111/1755-0998.13840

Mariani, S., Harper, L. R., Collins, R. A., Baillie, C., Wangensteen, O. S., McDevitt, A. D., … Genner, M. J. (2021). Estuarine molecular bycatch as a landscape-wide biomonitoring tool. Biological Conservation, 261, 109287. doi:10.1016/j.biocon.2021.109287

Martin, M. (2011). Cutadapt removes adapter sequences from high-throughput sequencing reads. EMBnet.Journal, 17(1), 10. doi:10.14806/ej.17.1.200

Mauvisseau, Q., Harper, L. R., Sander, M., Hanner, R. H., Kleyer, H., & Deiner, K. (2022, May 3). The Multiple States of Environmental DNA and What Is Known about Their Persistence in Aquatic Environments. Environmental Science and Technology. American Chemical Society. doi:10.1021/acs.est.1c07638

McClenaghan, B., Fahner, N., Cote, D., Chawarski, J., McCarthy, A., Rajabi, H., … Hajibabaei, M. (2020). Harnessing the power of eDNA metabarcoding for the detection of deep-sea fishes. PLOS ONE, 15(11), e0236540. doi:10.1371/journal.pone.0236540

Min, M. A., Barber, P. H., & Gold, Z. (2021). MiSebastes: An eDNA metabarcoding primer set for rockfishes (genus Sebastes). Conservation Genetics Resources, 13(4), 447–456. doi:10.1007/s12686-021-01219-2

Miya, M., Sato, Y., Fukunaga, T., Sado, T., Poulsen, J. Y., Sato, K., … Iwasaki, W. (2015). MiFish, a set of universal PCR primers for metabarcoding environmental DNA from fishes: Detection of more than 230 subtropical marine species. Royal Society Open Science, 2(7). doi:10.1098/rsos.150088

Newton, J. P., Bateman, P. W., Heydenrych, M. J., Kestel, J. H., Dixon, K. W., Prendergast, K. S., … Nevill, P. (2023). Monitoring the birds and the bees: Environmental <scp>DNA</scp> metabarcoding of flowers detects plant–animal interactions. Environmental DNA, 5(3), 488–502. doi:10.1002/edn3.399

Piñol, J., Mir, G., Gomez-Polo, P., & Agustí, N. (2015). Universal and blocking primer mismatches limit the use of high-throughput DNA sequencing for the quantitative metabarcoding of arthropods. Molecular Ecology Resources, 15(4), 819–830. doi:10.1111/1755-0998.12355

Piñol, Josep, Senar, M. A., & Symondson, W. O. C. (2019). The choice of universal primers and the characteristics of the species mixture determine when DNA metabarcoding can be quantitative. Molecular Ecology, 28(2). doi:10.1111/mec.14776

R Core Team. (2022). R: A Language and Environment for Statistical Computing. Vienna, Austria: R Foundation for Statistical Computing. Retrieved from https://www.r-project.org/

Riaz, T., Shehzad, W., Viari, A., Pompanon, F., Taberlet, P., & Coissac, E. (2011). EcoPrimers: Inference of new DNA barcode markers from whole genome sequence analysis. Nucleic Acids Research, 39(21), e145–e145. doi:10.1093/nar/gkr732

Ritter, C. D., Dunthorn, M., Anslan, S., de Lima, V. X., Tedersoo, L., Nilsson, R. H., & Antonelli, A. (2020). Advancing biodiversity assessments with environmental DNA: Long-read technologies help reveal the drivers of Amazonian fungal diversity. Ecology and Evolution, 10(14), 7509–7524. doi:10.1002/ece3.6477

Sales, N. G., McKenzie, M. B., Drake, J., Harper, L. R., Browett, S. S., Coscia, I., … McDevitt, A. D. (2020). Fishing for mammals: Landscape-level monitoring of terrestrial and semi-aquatic communities using eDNA from riverine systems. Journal of Applied Ecology, 57(4), 707– 716. doi:10.1111/1365-2664.13592

Stat, M., Huggett, M. J., Bernasconi, R., Dibattista, J. D., Berry, T. E., Newman, S. J., … Bunce, M. (2017). Ecosystem biomonitoring with eDNA: Metabarcoding across the tree of life in a tropical marine environment. Scientific Reports, 7(1), 1–11. doi:10.1038/s41598-017-12501-5

Taberlet, P., Bonin, A., Zinger, L., & Coissac, E. (2018). Environmental DNA: For biodiversity research and monitoring. Environmental DNA: For Biodiversity Research and Monitoring. Oxford University Press. doi:10.1093/oso/9780198767220.001.0001

Takahashi, M., Saccò, M., Kestel, J. H., Nester, G., Campbell, M. A., van der Heyde, M., … Allentoft, M. E. (2023, May 15). Aquatic environmental DNA: A review of the macro-organismal biomonitoring revolution. Science of the Total Environment. Elsevier B.V. doi:10.1016/j.scitotenv.2023.162322

Thalinger, B., Deiner, K., Harper, L. R., Rees, H. C., Blackman, R. C., Sint, D., … Bruce, K. (2021). A validation scale to determine the readiness of environmental DNA assays for routine species monitoring. Environmental DNA, 00, 1–14. https://doi.org/10.1002/edn3.189

Thalinger, B., Kirschner, D., Pütz, Y., Moritz, C., Schwarzenberger, R., Wanzenböck, J., & Traugott, M. (2021). Lateral and longitudinal fish environmental DNA distribution in dynamic riverine habitats. Environmental DNA, 3(1), 305–318. doi:10.1002/edn3.171

Tournayre, O., Leuchtmann, M., Filippi-Codaccioni, O., Trillat, M., Piry, S., Pontier, D., … Galan, M. (2020). In silico and empirical evaluation of twelve metabarcoding primer sets for insectivorous diet analyses. Ecology and Evolution, 10(13), 6310–6332. doi:10.1002/ece3.6362

Tsuji, S., Takahara, T., Doi, H., Shibata, N., & Yamanaka, H. (2019). The detection of aquatic macroorganisms using environmental DNA analysis—A review of methods for collection, extraction, and detection. Environmental DNA, 1(2), 99–108. doi:10.1002/edn3.21

Untergasser, A., Cutcutache, I., Koressaar, T., Ye, J., Faircloth, B. C., Remm, M., & Rozen, S. G. (2012). Primer3-new capabilities and interfaces. Nucleic Acids Research, 40(15), e115– e115. doi:10.1093/nar/gks596

Ushio, M., Murata, K., Sado, T., Nishiumi, I., Takeshita, M., Iwasaki, W., & Miya, M. (2018). Demonstration of the potential of environmental DNA as a tool for the detection of avian species. Scientific Reports, 8(1), 4493. doi:10.1038/s41598-018-22817-5

Valsecchi, E., Bylemans, J., Goodman, S. J., Lombardi, R., Carr, I., Castellano, L., … Galli, P. (2020). Novel universal primers for metabarcoding environmental DNA surveys of marine mammals and other marine vertebrates. Environmental DNA, 2(4), 460–476. doi:10.1002/edn3.72

Weigand, H., Beermann, A. J., Čiampor, F., Costa, F. O., Csabai, Z., Duarte, S., … Ekrem, T. (2019, August 15). DNA barcode reference libraries for the monitoring of aquatic biota in Europe: Gap-analysis and recommendations for future work. Science of the Total Environment. Elsevier B.V. doi:10.1016/j.scitotenv.2019.04.247

Wickham, H. (2016). ggplot2: Elegant Graphics for Data Analysis. Springer-Verlag New York. Retrieved from https://ggplot2.tidyverse.org

Wickham, Hadley, François, R., Henry, L., & Müller, K. (2019). dplyr: A Grammar of Data Manipulation. Retrieved from https://cran.r-project.org/package=dplyr

Yang, J., Zhang, L., Mu, Y., & Zhang, X. (2023). Small changes make big progress: A more efficient <scp>eDNA</scp> monitoring method for freshwater fish. Environmental DNA, 5(2), 363–374. doi:10.1002/edn3.387

Yates, M. C., Furlan, E., Thalinger, B., Yamanaka, H., & Bernatchez, L. (2023). Beyond species detection **—** leveraging environmental DNA and environmental RNA to push beyond presence/absence applications. Environmental DNA. doi:10.1002/edn3.459

Yu, D. W., Ji, Y., Emerson, B. C., Wang, X., Ye, C., Yang, C., & Ding, Z. (2012). Biodiversity soup: metabarcoding of arthropods for rapid biodiversity assessment and biomonitoring. Methods in Ecology and Evolution, 3(4), 613–623. doi:10.1111/j.2041-210X.2012.00198.x

Zhang, S., Zhao, J., & Yao, M. (2020). A comprehensive and comparative evaluation of primers for metabarcoding eDNA from fish. Methods in Ecology and Evolution, 11(12), 1609–1625. doi:10.1111/2041-210X.13485

Zhang, S., Zhao, J., & Yao, M. (2023). Urban landscape-level biodiversity assessments of aquatic and terrestrial vertebrates by environmental DNA metabarcoding. Journal of Environmental Management, 340, 117971. doi:10.1016/j.jenvman.2023.117971

