## Supporting Information 5 for "BirT: a novel primer pair for avian environmental DNA metabarcoding"

SI5: Optimization of PCR conditions

To optimize the annealing temperature in the first PCR (Elbrecht & Steinke, 2018) for bird eDNA amplification, we carried out tests with different types of target and non-target extracts. As target DNA extracts we selected i) three bird tissue extracts (*Ara chloropterus, Branta canadensis, Phalacrocorax auritus*), , ii) two DNA extracts of mixed bird feathers collected from nests around Guelph, Canada (Levesque-Beaudin, Steinke, Böcker, & Thalinger, 2023), and iii) two extracts per filter pore size of the Smith Root filters and three extracts obtained from the NatureMetrics filters. The non-target extracts, which were highly unlikely to contain bird DNA, were two horse fecal samples and two fish tissues (*Salmo salar, Oncorhynchus mykiss*; Table SI4.1).

**Table SI5.1:** The target and non-target DNA extracts used to optimize cycle number and annealing temperature in the first of two consecutive PCRs for metabarcoding with the BirT primers. Additionally, the position of each extract in the Gel-Pictures (Fig. SI4.1) is indicated.

| sample type | sample name | DNA concentration (ng/µl) | Gel position (no spike-in) | Gel position with spike-in |
| --- | --- | --- | --- | --- |
| target | *Ara chloropterus* 1 high conc. |  | 1 |  |
| target | *Branta canadensis* 1 high conc. |  | 2, 25 |  |
| target | *Phalacrocorax auritus* 1 high conc. |  | 3 |  |
| target | *Ara chloropterus* 1 low conc. |  | 4 |  |
| target | *Branta canadensis* 1 low conc. |  | 5 |  |
| target | *Phalacrocorax auritus* 1 low conc. |  | 6 |  |
| target | mixed feather extract from nest (f3) |  | 7 |  |
| target | mixed feather extract from nest (f7) |  | 8 |  |
| target | Smith Root filter 1.2µm (MB 8) | 10.60 | 17, 39 |  |
| target | Smith Root filter 1.2µm (MB17) | 29.40 | 18 |  |
| target | Smith Root filter 5µm (MB6) | 1.51 | 19 |  |
| target | Smith Root filter 5µm (MB15) | 8.62 | 20 |  |
| target | NatureMetrics filter (MB1) | 25.20 | 21 |  |
| target | NatureMetrics filter (MB4) | 13.20 | 22 |  |
| target | NatureMetrics filter (MB19) | 7.17 | 23, 40 |  |
| non-target | horse feces (DR1) |  | 9, 26, 28 | 11, 29, 31 |
| non-target | horse feces (DR2) |  | 10, 27 | 12, 30 |
| non-target | *Salmo salar* 1 | 1.00 | 13, 33 | 15, 35, 37 |
| non-target | *Oncorhynchus mykiss* 1 | 1.00 | 14, 34 | 16, 36, 38 |
| negative control | PCR-grade water |  | 24, 32 |  |

All target and non-target extracts were run in 20 µl of total volume containing 10 µl of 2 × Multiplex Master Mix (Qiagen), 1 µl of the BirT-F and BirT-R primers (10 µM), respectively, 2 µl DNA extract, and molecular grade water. Additionally, 2 µl of the high-concentration *B. canadensis* extract were added (instead of molecular grade water) to wells containing a horse feces extract and the *Salmo salar* extract. This was repeated for the low-concentration version of the *B. canadensis* extract, leading to a total of 18 wells including one negative control.

These PCR mixes were run in a gradient PCR with thermocycling conditions as follows: 10 min of initial denaturation at 95°C, 25 cycles of denaturation at 95°C for 30 s, annealing at either 58, 60, or 62 °C for 30 s, elongation at 72°C for 1 min, and final extension at 72°C for 10 min, once. The PCR war repeated with the exact same setup but 35 cycles instead of 25. All PCR products were subjected to a second PCR using 20 µl reactions with 10 µl of 2 × Multiplex Master Mix, 1 µl of a forward and a reverse fusion primer (10 µM; including Illumina sequencing adapters), 2 µl of PCR product, and molecular grade water. The thermocycling conditions were the same as for the first PCR, except for an annealing temperature of 50 °C and the consistent use of 25 cycles.

After the second PCR, amplification success was examined on 1.5% agarose gels. Optimum amplification conditions were defined as i) no amplification of non-target extracts, ii) strong amplification of target extracts and non-target extracts spiked with *B. canadensis*, iii) minimal non-target bands such as primer dimers. This led to the selection of 25 cycles and 60 °C annealing temperature as optimum conditions in the first PCR for the screening of field-collected bird eDNA samples.

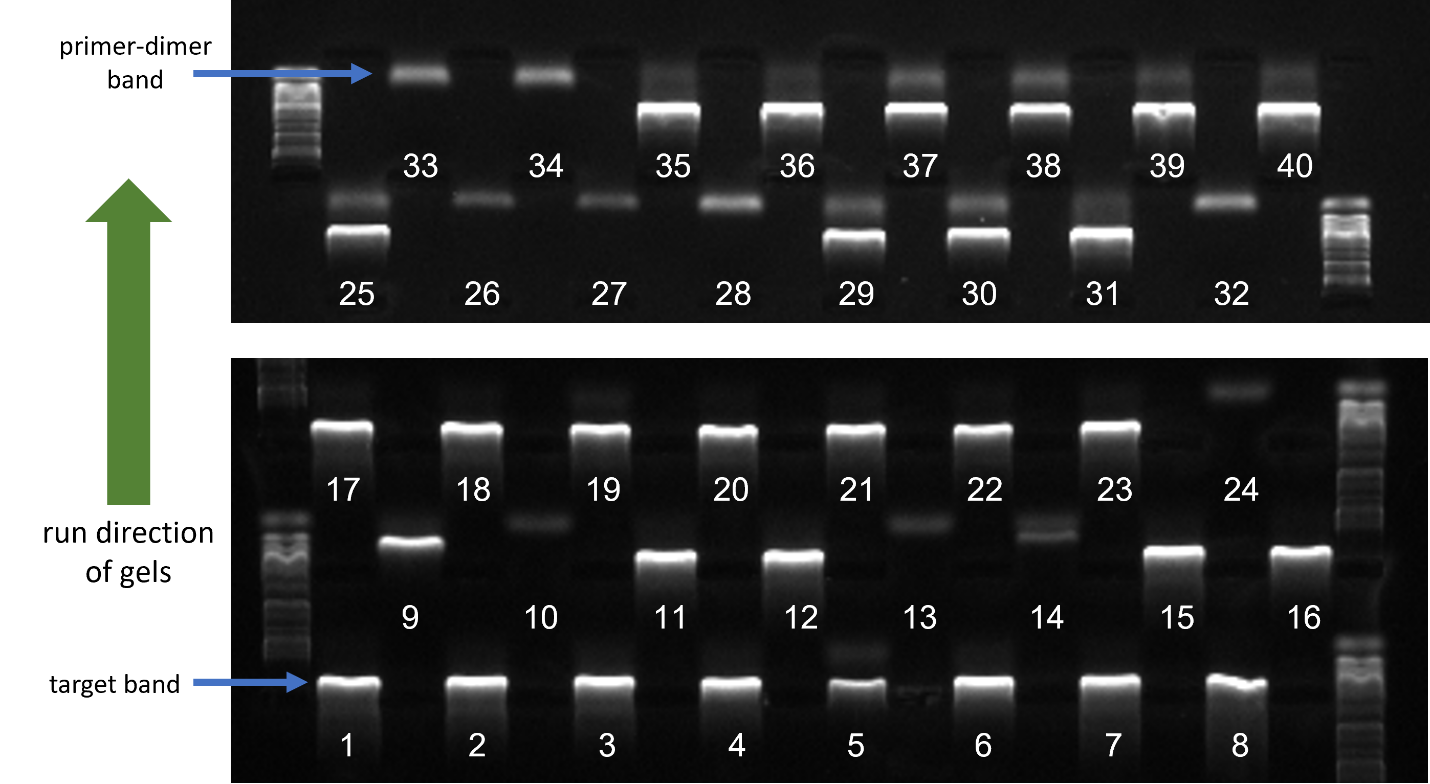

**Figure SI5.1:** Gel pictures showing the amplification after the second round of PCR for the selected optimum PCR conditions. The origin of each band is indicated in Table SI4.1. Please note, that the bands produced in the absence of target DNA (e.g. in positions 9 or 33) are generated by primer-dimers.

**References**

Elbrecht, V., & Steinke, D. (2018). Scaling up DNA metabarcoding for freshwater macrozoobenthos monitoring. *Freshwater Biology*, *64*(2), fwb.13220. doi:10.1111/fwb.13220

Levesque-Beaudin, V., Steinke, D., Böcker, M., & Thalinger, B. (2023). Unravelling bird nest arthropod community structure using metabarcoding. *BioRxiv*, 2023.03.09.531929. doi:10.1101/2023.03.09.531929
